## Supplementary material for "Distinctive roles of translesion polymerases DinB1 and DnaE2 in diversification of the mycobacterial genome through substitution and frameshift mutagenesis": SI Figs and Tables

**Figure S1. *dinB1* expression phenotypes in WT, Δ*dnaE2* and Δ*recA* backgrounds.** (A) Growth of *M. smegmatis* carrying the *dinB1*$^{Msm}$ expression plasmid on agar medium containing the indicated concentrations of inducer in agar. (B) Liquid growth of *M. smegmatis* carrying the *dinB1*$^{Msm}$ expression plasmid in presence of the indicated concentrations of inducer. (C) Liquid growth of the indicated strains in presence of inducer (50nM). (D) Rif$^{R}$ frequency in indicated strains in presence of inducer. Results shown are means (± SEM) of data obtained from biological replicates symbolized by grey dots. Stars above the means mark a statistical difference with the reference strain (empty vector or 0nM of inductor) (**, $P<0.01$; ***, $P<0.001$). (E) CLUSTAL O (1.2.4) multiple sequence alignment of *E. coli* (Eco), *M. smegmatis* (Msm) and *M. tuberculosis* (Mtb) DinB1 sequences.

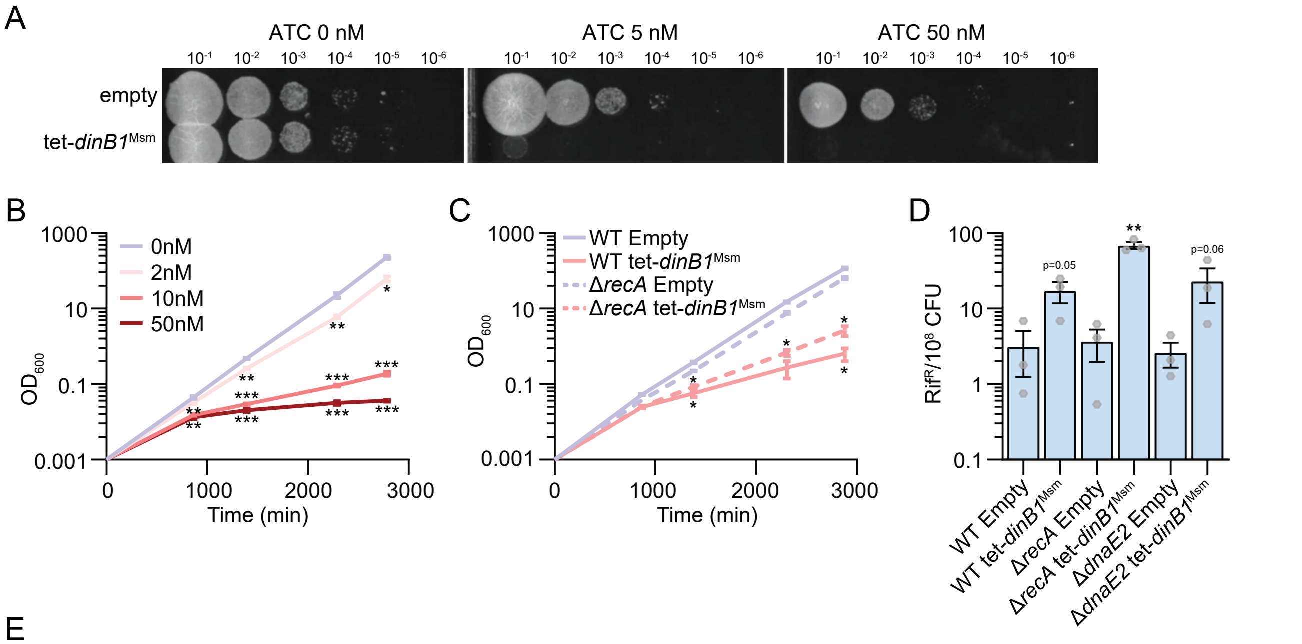


E

```
Eco  ----------MRKIIHVDMDCFFAAVEMRDNPALRDIPIAIGGSRERRGVISTANYPARK 50
Msm  MEGTVARTASRRWVLHLDMDAFFASVEQLTRPTLRGRPVLVGGLG-GRGVVAGASYEARR 59
Mtb  --------VESRWVLHLDMDAFFASVEQLTRPTLRGRPVLVGGLG-GRGVVAGASYEARA 51
                 *  ::*:***.***:**    .*:**.  *:  :**     ***::  *.*  **

Eco  FGVRSAMPTGMALKLCPH-LTLLPGRFDAYKEASNHIREIFSRYTSRIEPLSLDEAYLDV 109
Msm  YGARSAMPMHQARRLVGAPAVVLPPRGAVYGVASRRALDTVRSVVPVLEQLSFDEAFGEP 119
Mtb  YGARSAMPMHQARRLIGVTAVVLPPRGVVYGIASRRVFDTVRGLVPVVEQLSFDEAFAEP 111
     :*.*****    *  :*        .:** *   .*   **.:   :  .     .   :*  **:***: :

Eco  TDSVHCH-GSATLIAQEIRQTIFNELQLTASAGVAPVKFLAKIASDMNKPNGQFVITPAE 168
Msm  SELAGAEAADVEAFCERLRAKVLEHTGLVASVGAGSGKQIAKIASGLAKPDGIRVVRREE 179
Mtb  PQLAGAVAEDVETFCERLRRRVRDETGLIASVGAGSGKQIAKIASGLAKPDGIRVVRHAE 171
      :  .  .    ..   :.:.:*   :  :.    *  **.*..   *  :*****.:  **:*   *:    *

Eco  VPAFLQTLPLAKIPGVGKVSAAKLEAMGLRTCGDVQKCDLVMLLKRF-GKFGRILWERSQ 227
Msm  EAVLLHGLPVRKLWGIGPVAEDRLHRIGIETIGAFAALTEAEAANVLGSTIGPALHRLAR 239
Mtb  EQALLSGLPVRRLWGIGPVAEEKLHRLGIETIGQLAALSDAEAANILGATIGPALHRLAR 231
       .:*   **:  ::  *:*  *:   :*.  :*:.*  *  .       .    :  :  ..:*   *  .  ::

Eco  GIDERDVNSERLRKSVGVERTMAEDIHHWSECEAIIERLYPELERRLAKVKPDLLIARQG 287
Msm  GIDDRPVAERAEAKQISAESTFPEDLTTLAQLQDAIVPIGEHAHRRLEKDGRGA--RTVT 297
Mtb  GIDDRPVVERAEAKQISAESTFAVDLTTMEQLHEAIDSIAEHAHQRLLRDGRGA--RTIT 289
     ***:*  *  ..    *.:..*  *:   *:      :  .   *   :   .  .:**  :    .

Eco  VKLKFDDFQQTTQE--HVWPRLNKADLIATARKTWDERRGGRGVRLVGLHVTLLDPQMER 345
Msm  VKLKKSDMSTLTRSATLAYATTEASTLIGTARRLLLDPVEIGPIRLVGVGFSGLSDIRQE 357
Mtb  VKLKKSDMSTLTRSATMPYPTTDAGALFTVARRLLPDPLQIGPIRLLGVGFSGLSDIRQE 349
     **** .*:.   *:.      :     :  .  *:  .**:     :        :**:*:  .:  *.     :.

Eco  QLVLGL------------------------------------------------------ 351
Msm  SLFPDLEQPEEFPDAAPQ-VESVQTGPSSPTTQWRVGDDVAHTELGHGWVQGAGHGVMTV 416
Mtb  SLFADSDLTQETAAAHYVETPGAVVPAAHDATMWRVGDDVAHPELGHGWVQGAGHGVVTV 409
     .*.  .
Eco  ------------------------------------------------------------ 351
Msm  RFETRASGPGPARTFPEDSPEISRADPVDSLDWAEYLSSLADYQSTP------------ 463
Mtb  RFETRGSGPGSARTFPVDTGDISNASPLDSLDWPDYIGQLSVEGSAGASAPTVDDVGDR 468
```

DinB1[Mtb] extra N-ter 5aa

Metal-binding aspartate residues

β-clamp binding domain

| | Eco | Msm | Mtb |
|---|---|---|---|
| Eco | 100.00 | 30.17 | 29.60 |
| Msm | 30.17 | 100.00 | 74.73 |
| Mtb | 29.60 | 74.73 | 100.00 |

Figure S1. *dinB1* expression phenotypes in WT, Δ*dnaE2* and Δ*recA* backgrounds.

**Figure S2. Expression levels of mycobacterial TLS polymerases.** Expression level of the indicated *M. smegmatis* genes in indicated genetic backgrounds and conditions measured by RNA sequencing or RT-qPCR. Results shown are means (± SEM) of data obtained from biological replicates symbolized by grey dots. Stars above bars mark a statistical difference with the reference strain (no treatment in the same strain) (*, P<0.05; **, P<0.01; ***, P<0.001).

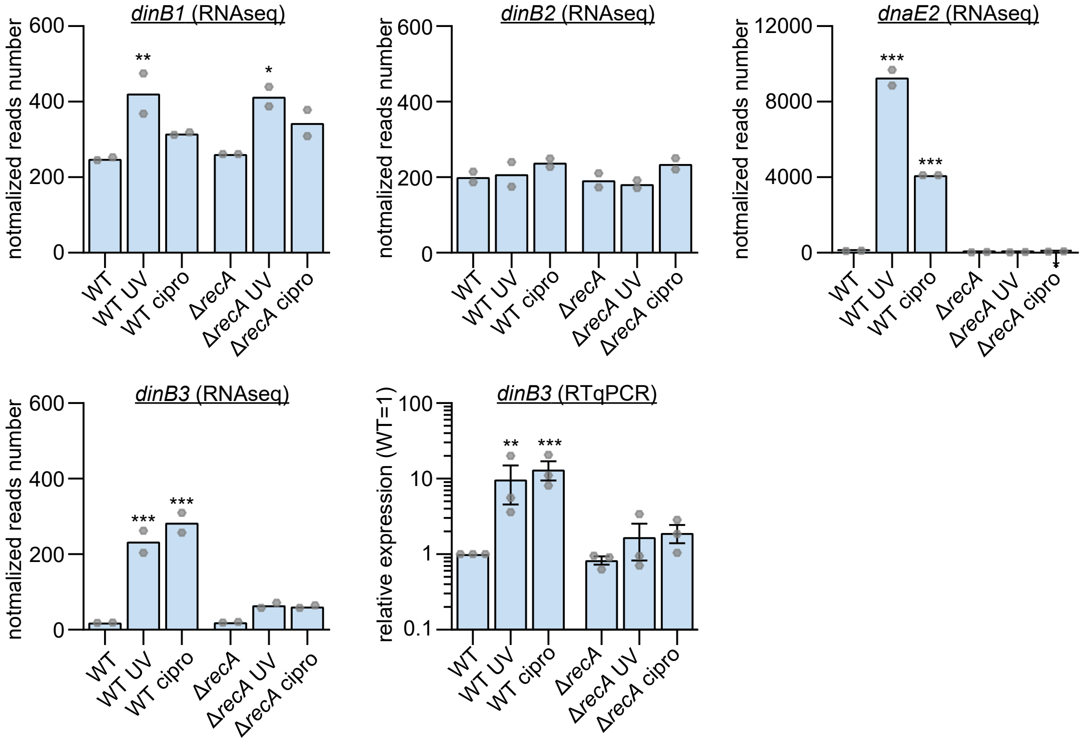


Figure S2. Expression level of mycobacterial TLS polymerases.

**Figure S3. Role of TLS polymerases in DNA damage tolerance.** (A) $H_2O_2$, (B) ciprofloxacin, (D) MMC or (E) 4-NQO sensitivity of indicated strains measured by disc diffusion assay. (C) Viability of indicated strains after treatment with indicated doses of UV. Results shown are means (± SEM) of data obtained from biological replicates symbolized by grey dots. Stars above the means mark a statistical difference with the reference strain (WT) (**, $P<0.01$; ***, $P<0.001$). (F) Growth of indicated strains on agar medium containing the indicated concentrations of MMS in agar. (G) Pictures of disc diffusion assays with indicated strains and performed to measure the sensitivity of *M. smegmatis* to the indicated chemical agents.

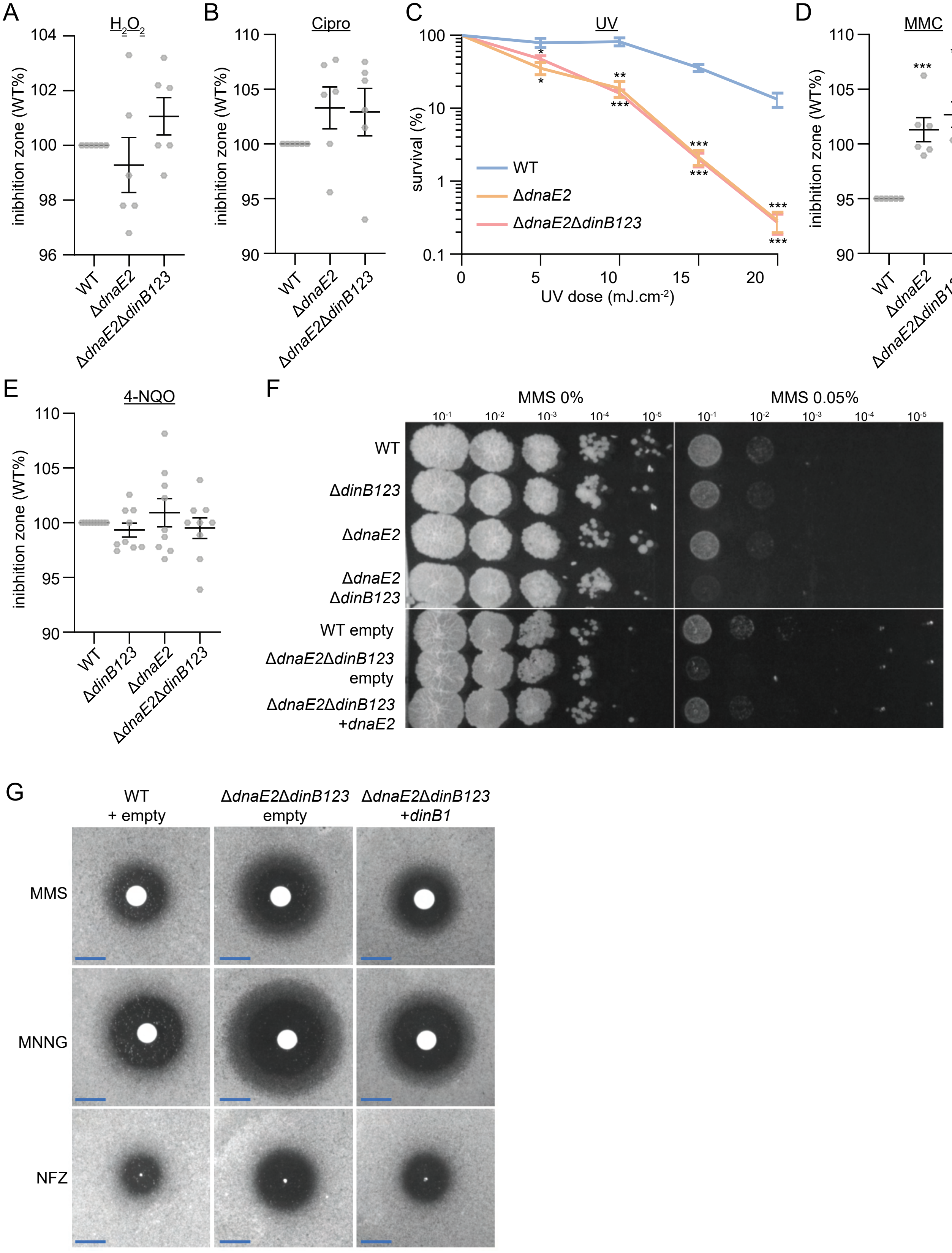


Figure S3. Role of TLS polymerases in DNA damage tolerance.

**Figure S4. -1 and +1 frameshift mutagenesis in diverse homo-oligonucleotide runs after expression of *dinB1*.** (A) leu+ or (B, C, D, E and F) $kan^R$ frequency in indicated strains in presence of inducer and relative (pie chart) or absolute (bar chart) frequencies of nucleotide changes detected in *leuD* or *kan* coded by color. The number of sequenced $leu^+$ or $kan^R$ colonies is given in the center of each pie chart. The sequence in (A) shows the *leuD*$^{-1}$ reporter designed to measure +1 frameshift (FS) with the N termini of *leuD* and the C termini of the upstream *leuC*. The blue box shows the 1 base pair deletion introduced to inactivate *leuD* and the green boxes show the homonucleotide runs in which the majority of -1 FS have been detected in $leu^+$ cells. Results shown are means (± SEM) of data obtained from biological replicates symbolized by grey dots. Stars above bars mark a statistical difference with the reference strain (empty) (*, P<0.05; **, P<0.01; ***, P<0.001).

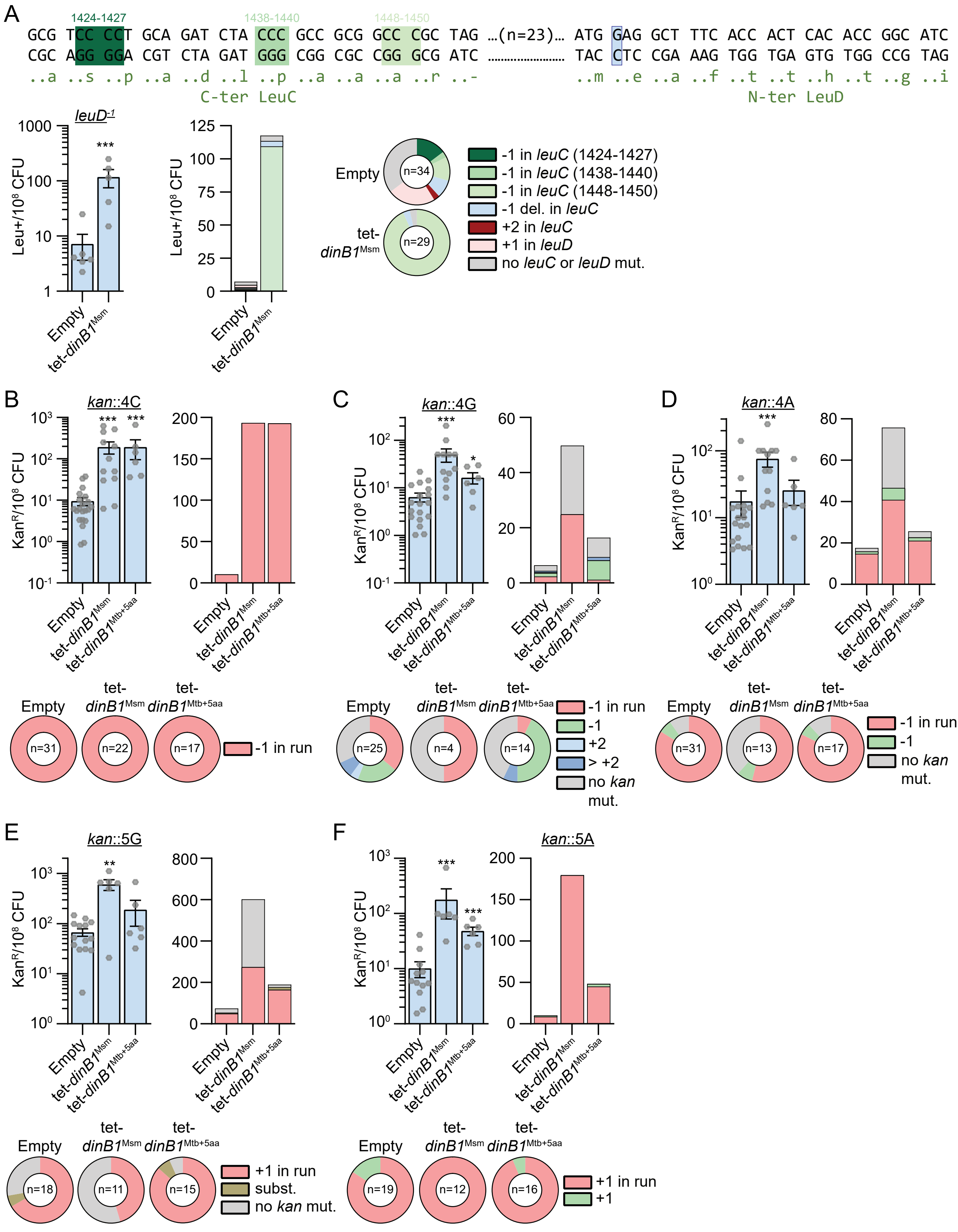


Figure S4. -1 and +1 frameshift mutagenesis in diverse homonucleotide runs after expression of *dinB1*.

**Figure S5. DinB1 discriminates against ddNTPs**. Reaction mixtures containing 10 mM Tris-HCl, pH 7.5, 5 mM $MnCl_2$,1 pmol 5' $^{32}P$-labeled primer-template DNAs with A6 or T6 runs in the template strand (depicted below, and included as indicated above the lanes), 125 μM nucleotides as specified, and 10 pmol DinB1 were incubated at 37˚C for 15 min. DinB1 was omitted from control reactions in lanes –. The reaction products were analyzed by urea-PAGE and visualized by autoradiography. "Faithful" +6 and +9 primer extension products are denoted by arrowheads.

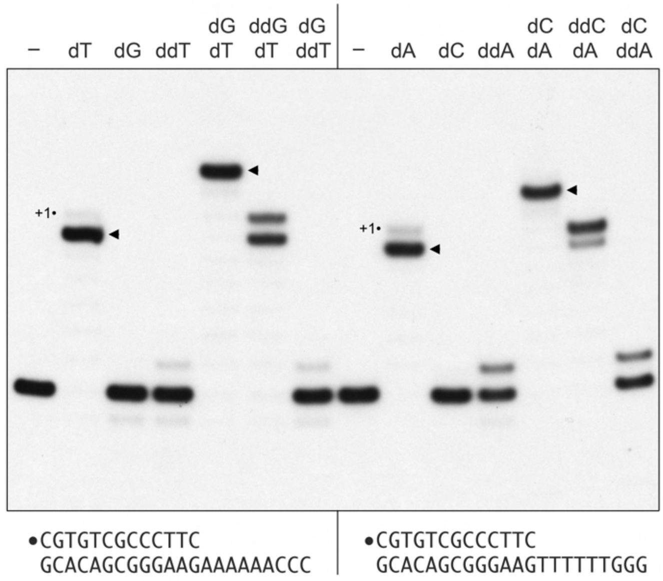


Figure S5. DinB1 discriminates against ddNTPs.

**Figure S6. Spontaneous -1 and +1 frameshift mutagenesis in diverse homonucleotide runs.** $Kan^R$ frequency in indicated strains and relative (pie chart) or absolute (bar chart) frequencies of nucleotide changes detected in *kan* coded by color. The number of sequenced $kan^R$ colonies is given in the center of each pie chart. Results shown are means (± SEM) of data obtained from biological replicates symbolized by grey dots. Stars above bars mark a statistical difference with the reference strain (WT) (*, $P<0.05$).

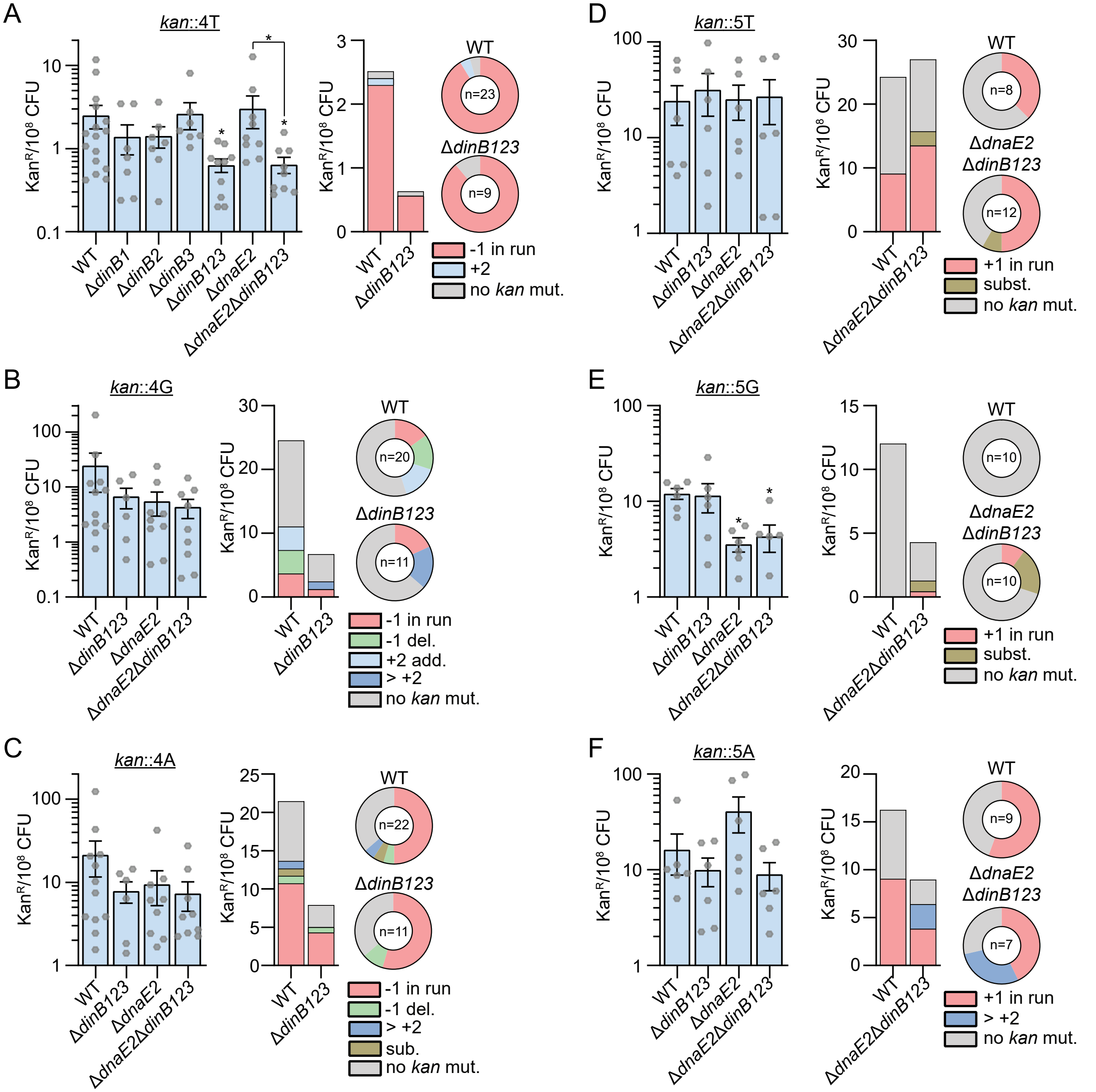


Figure S6. Spontaneous -1 and +1 frameshift mutagenesis in diverse homonucleotide runs.

**Figure S7. UV-induced -1 and +1 frameshift mutagenesis in diverse homonucleotide runs.** Kan$^R$ frequency in indicated strains without (blue) or with UV treatment (green) and relative (pie chart) or absolute (bar chart) frequencies of nucleotide changes detected in *kan* coded by color. The number of sequenced kan$^R$ colonies is given in the center of each pie chart. Results shown are means (± SEM) of data obtained from biological replicates symbolized by grey dots. Stars above bars mark a statistical difference with the reference strain (WT+UV) and lines connecting two strains show a statistical difference between them (*, $P<0.05$; **, $P<0.01$; ***, $P<0.001$).

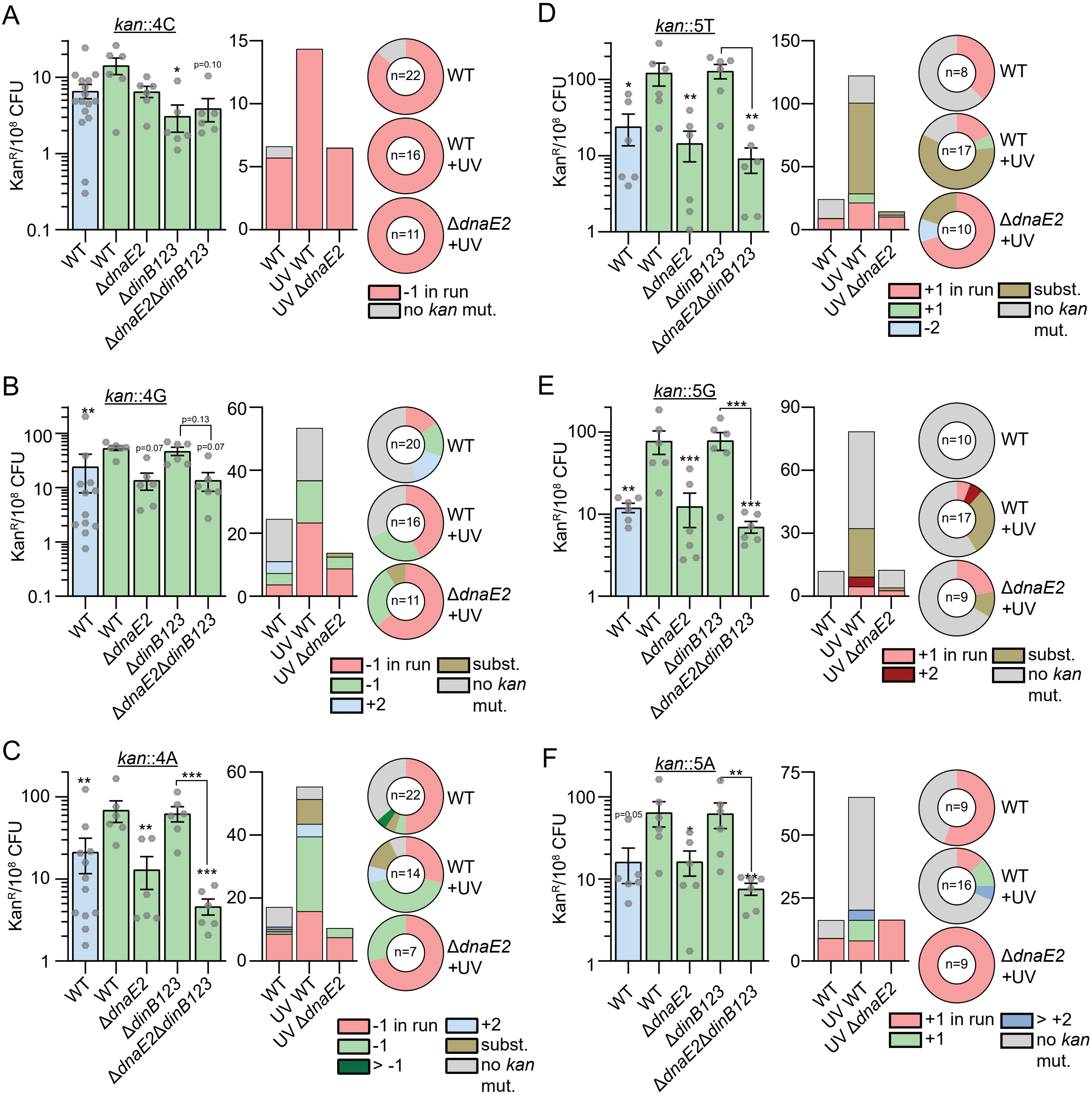


Figure S7. UV-induced -1 and +1 frameshift mutagenesis in diverse homonucleotide runs.

### Supplementary material tables

Table S1: plasmids used in this work and cloning methods

| Plasmids | description | Cloning enzyme sites | Cloning primers (infusion reaction) | references or sources |
|---|---|---|---|---|
| pmsg419 | Empty ATc-on system vector ($Hyg^R$, OriMyc) | | | Lab Stock |
| pAJF067 | Gene replacement vector ($Hyg^R$, *galK*, *sacB*) | | | Fay and Glickman, 2014 |
| pDB60 | Complementation vector ($Strep^R$, attP(L5)) | | | Lab Stock |
| pDB60-*dnaE2* | pDB60 derivative for *dnaE2*$^{Msm}$ complementation | *BstB1* | OAM118-OAM185 | This work |
| pDP64 | pmsg419 derivative for *dinB1*$^{Msm}$ overexpression | *Cla1* | ODP290-ODP291 | This work |
| pDP65 | pmsg419 derivative for *dinB1*$^{ST}$overexpression | *Cla1* | ODP290-ODP292 | This work |
| pDP66 | pmsg419 derivative for *dinB1*$^{D113A\text{-}ST}$ | *Cla1* | ODP290-ODP293+ODP294-ODP292 | This work |
| pDP67 | pmsg419 derivative for *dinB1*$^{F23L\text{-}ST}$ | *Cla1* | ODP290-ODP295+ODP296-ODP292 | This work |
| pDP88 | pmsg419 derivative for *dinB1*$^{Mtb}$ overexpression | *Cla1* | ODP343-ODP344 | This work |
| pDP104 | pAJF067 derivative for *leuD*$^{\Delta4}$ deletion (*leuD*$^{-1}$) | *Nde1* | ODP389-ODP390+ODP391-ODP392 | This work |
| pDP105 | pAJF067 derivative for *leuD*$^{\Delta4\text{-}5}$ deletion (*leuD*$^{-2}$) | *Nde1* | ODP389-ODP390+ODP393-ODP392 | This work |
| pDP112 | pDB60 derivative for *dinB1*$^{Msm}$ complementation | *EcoR1* | ODP480-ODP481 | This work |
| pDP114 | pDB60 derivative for *dinB2*$^{Msm}$ complementation | *EcoR1* | ODP425-ODP426+ODP427-ODP428 | This work |
| pDP115 | pDB60 derivative for *dinB3*$^{Msm}$ complementation | *EcoR1* | ODP429-ODP430 | This work |
| pDP117 | pmsg419 derivative for *dinB1*$^{\Delta\beta clamp}$ overexpression | *Cla1* | ODP290-ODP433+ODP434-ODP291 | This work |
| pDP118 | pDB60 derivative for *dinB1*$^{Mtb}$ complementation | *EcoR1* | ODP435-ODP436 | This work |
| pDP119 | pDB60 derivative for *dinB2*$^{Mtb}$ complementation | *EcoR1* | ODP437-ODP438 | This work |
| pDP120 | pDB60 derivative with *kan*::3T | *EcoR1* | ODP443-ODP445+ODP446-ODP444 | This work |
| pDP121 | pDB60 derivative with *kan*::3C | *EcoR1* | ODP443-ODP445+ODP447-ODP444 | This work |
| pDP122 | pDB60 derivative with *kan*::3G | *EcoR1* | ODP443-ODP445+ODP448-ODP444 | This work |
| pDP123 | pDB60 derivative with *kan*::3A | *EcoR1* | ODP443-ODP445+ODP449-ODP444 | This work |
| pDP124 | pDB60 derivative with *kan*::4T | *EcoR1* | ODP443-ODP445+ODP450-ODP444 | This work |
| pDP125 | pDB60 derivative with *kan*::4C | *EcoR1* | ODP443-ODP445+ODP451-ODP444 | This work |
| pDP126 | pDB60 derivative with *kan*::4G | *EcoR1* | ODP443-ODP445+ODP452-ODP444 | This work |
| pDP127 | pDB60 derivative with *kan*::4A | *EcoR1* | ODP443-ODP445+ODP453-ODP444 | This work |
| pDP128 | pDB60 derivative with *kan*::6T | *EcoR1* | ODP443-ODP445+ODP454-ODP444 | This work |
| pDP129 | pDB60 derivative with *kan*::6C | *EcoR1* | ODP443-ODP445+ODP455-ODP444 | This work |
| pDP130 | pDB60 derivative with *kan*::6G | *EcoR1* | ODP443-ODP445+ODP456-ODP444 | This work |
| pDP131 | pDB60 derivative with *kan*::6A | *EcoR1* | ODP443-ODP445+ODP457-ODP444 | This work |
| pDP144 | pDB60 derivative with *kan*::5T | *EcoR1* | ODP443-ODP445+ODP490-ODP444 | This work |
| pDP145 | pDB60 derivative with *kan*::5C | *EcoR1* | ODP443-ODP445+ODP491-ODP444 | This work |
| pDP146 | pDB60 derivative with *kan*::5G | *EcoR1* | ODP443-ODP445+ODP492-ODP444 | This work |
| pDP147 | pDB60 derivative with *kan*::5A | *EcoR1* | ODP443-ODP445+ODP493-ODP444 | This work |
| pDP157 | pmsg419 derivative for *dinB1*$^{Mtb+5aa}$ overexpression | *Cla1* | ODP514-ODP344 | This work |

Table S2: *rpoB* mutations incorporated by TLS polymerases

| | Empty | | *dinB1*$^{Msm}$ OE | | *dinB1*$^{Mtb+5aa}$ OE | | Δ*dnaE2* $H_2O_2$ | | WT $H_2O_2$ | |
|---|---|---|---|---|---|---|---|---|---|---|
| mutations | prop. in % | mut./$10^8$ | prop. in % | mut./$10^8$ | prop. in % | mut./$10^8$ | prop. in % | mut./$10^8$ | prop. in % | mut./$10^8$ |
| Leu427(CTG>CCT) | 0.00 | 0.00 | 0.00 | 0.00 | 0.00 | 0.00 | 6.82 | 1.30 | 4.00 | 3.00 |
| Ser428(TCG>TGG) | 0.00 | 0.00 | 0.00 | 0.00 | 0.00 | 0.00 | 9.09 | 1.73 | 2.00 | 1.50 |
| Gln429(CAG>AAG) | 2.17 | 0.12 | 0.00 | 0.00 | 4.17 | 1.84 | 2.27 | 0.43 | 2.00 | 1.50 |
| Gln429(CAG>CTG) | 0.00 | 0.00 | 2.22 | 0.76 | 0.00 | 0.00 | 0.00 | 0.00 | 4.00 | 3.00 |
| Gln429(CAG>CCG) | 1.09 | 0.06 | 0.00 | 0.00 | 0.00 | 0.00 | 0.00 | 0.00 | 0.00 | 0.00 |
| Asp432(GAC>TAC) | 2.17 | 0.12 | 0.00 | 0.00 | 0.00 | 0.00 | 2.27 | 0.43 | 0.00 | 0.00 |
| Asp432(GAC>AAC) | 0.00 | 0.00 | 0.00 | 0.00 | 0.00 | 0.00 | 2.27 | 0.43 | 0.00 | 0.00 |
| Asp432(GAC>GTC) | 2.17 | 0.12 | 0.00 | 0.00 | 0.00 | 0.00 | 0.00 | 0.00 | 2.00 | 1.50 |
| Asp432(GAC>GGC) | 2.17 | 0.12 | 4.44 | 1.53 | 0.00 | 0.00 | 0.00 | 0.00 | 0.00 | 0.00 |
| Asp432(GAC>GAG) | 0.00 | 0.00 | 0.00 | 0.00 | 0.00 | 0.00 | 0.00 | 0.00 | 0.00 | 0.00 |
| Asp432(GAC>TTC) | 0.00 | 0.00 | 0.00 | 0.00 | 0.00 | 0.00 | 0.00 | 0.00 | 2.00 | 1.50 |
| Asn435(AAC>AAG) | 0.00 | 0.00 | 0.00 | 0.00 | 0.00 | 0.00 | 0.00 | 0.00 | 6.00 | 4.49 |
| Ser438(TCG>TTG) | 16.30 | 0.90 | 2.22 | 0.76 | 4.17 | 1.84 | 2.27 | 0.43 | 12.00 | 8.99 |
| Ser438(TCG>TGG) | 0.00 | 0.00 | 0.00 | 0.00 | 0.00 | 0.00 | 0.00 | 0.00 | 4.00 | 3.00 |
| His442(CAC>TAC) | 15.22 | 0.84 | 6.67 | 2.29 | 0.00 | 0.00 | 9.09 | 1.73 | 10.00 | 7.49 |
| His442(CAC>AAC) | 0.00 | 0.00 | 0.00 | 0.00 | 0.00 | 0.00 | 0.00 | 0.00 | 2.00 | 1.50 |
| His442(CAC>GAC) | 9.78 | 0.54 | 0.00 | 0.00 | 4.17 | 1.84 | 9.09 | 1.73 | 8.00 | 5.99 |
| His442(CAC>CCC) | 3.26 | 0.18 | 4.44 | 1.53 | 0.00 | 0.00 | 0.00 | 0.00 | 2.00 | 1.50 |
| His442(CAC>CGC) | 21.74 | 1.19 | 73.33 | 25.21 | 75.00 | 33.19 | 2.27 | 0.43 | 2.00 | 1.50 |
| His442(CAC>CCG) | 0.00 | 0.00 | 0.00 | 0.00 | 0.00 | 0.00 | 0.00 | 0.00 | 2.00 | 1.50 |
| Arg445(CGT>TGT) | 0.00 | 0.00 | 0.00 | 0.00 | 0.00 | 0.00 | 2.27 | 0.43 | 4.00 | 3.00 |
| Arg445(CGT>CTT) | 0.00 | 0.00 | 2.22 | 0.76 | 0.00 | 0.00 | 0.00 | 0.00 | 0.00 | 0.00 |
| Arg445(CGT>CCT) | 0.00 | 0.00 | 0.00 | 0.00 | 0.00 | 0.00 | 2.27 | 0.43 | 2.00 | 1.50 |
| Ser447(TCG>TTG) | 4.35 | 0.24 | 0.00 | 0.00 | 0.00 | 0.00 | 22.73 | 4.33 | 16.00 | 11.98 |
| Ser447(TCG>TGG) | 0.00 | 0.00 | 0.00 | 0.00 | 4.17 | 1.84 | 13.64 | 2.60 | 2.00 | 1.50 |
| Leu449(CTG>CCG) | 4.35 | 0.24 | 2.22 | 0.76 | 0.00 | 0.00 | 2.27 | 0.43 | 0.00 | 0.00 |
| Gly450(GGC>AGC) | 0.00 | 0.00 | 0.00 | 0.00 | 0.00 | 0.00 | 0.00 | 0.00 | 2.00 | 1.50 |
| Gly450(GGC>TGC) | 1.09 | 0.06 | 0.00 | 0.00 | 0.00 | 0.00 | 0.00 | 0.00 | 0.00 | 0.00 |
| Pro480(CCT>CTT) | 1.09 | 0.06 | 0.00 | 0.00 | 0.00 | 0.00 | 0.00 | 0.00 | 0.00 | 0.00 |
| Ile488(ATC>TTC) | 1.09 | 0.06 | 0.00 | 0.00 | 4.17 | 1.84 | 0.00 | 0.00 | 0.00 | 0.00 |
| Ile488(ATC>ATG) | 0.00 | 0.00 | 0.00 | 0.00 | 0.00 | 0.00 | 2.27 | 0.43 | 0.00 | 0.00 |
| Ser490(TCG>TTG) | 0.00 | 0.00 | 0.00 | 0.00 | 0.00 | 0.00 | 2.27 | 0.43 | 2.00 | 1.50 |
| del. | 1.09 | 0.06 | 0.00 | 0.00 | 0.00 | 0.00 | 0.00 | 0.00 | 0.00 | 0.00 |
| no *rpoB* mut. | 10.87 | 0.60 | 2.22 | 0.76 | 4.17 | 1.84 | 6.82 | 1.30 | 8.00 | 5.99 |
| total | 100.00 | 5.49 | 100.00 | 34.38 | 100.00 | 44.26 | 100.00 | 19.07 | 100.00 | 74.90 |

prop. in %: relative frequency of *rpoB* mutations found in indicated strains (also shown in Figures 2C and 3C)
mut./$10^8$: absolute frequency of *rpoB* mutations, expressed in number of mutations per $10^8$ CFU found in indicated strains (also shown in Figures 2D and 3C).

Table S3: *leuD* mutations detected in indicated strains

| *leuD*$^{-2}$ mut. | empty | tet-*dinB1*$^{Msm}$ | tet-*dinB1*$^{Mtb+5aa}$ | WT | Δ*dinB1* | Δ*dinB2* | Δ*dinB3* | Δ*dinB123* | Δ*dnaE2* | Δ*dnaE2*Δ*dinB123* | WT+empty | Δ*dinB1*+empty | Δ*dinB1*+*dinB1* | WT UV | Δ*dnaE2* UV | Δ*dinB123* UV | Δ*dinB123*Δ*dnaE2* UV |
|---|---|---|---|---|---|---|---|---|---|---|---|---|---|---|---|---|---|
| T del. (nuc. 9-11) | 18 | 20 | 11 | 33 | | 23 | 22 | 3 | 14 | 7 | 10 | 2 | 19 | 5 | 6 | 5 | 0 |
| A del. (nuc. 1) | | | | | | | | 1 | | | | | | | | | |
| T del. (nuc. 2) | | | | | | 1 | | | | | | 4 | | | | | |
| G del. (nuc. 3) | | | | 2 | 3 | 2 | | 1 | 1 | | | 1 | 1 | | | 1 | |
| C del. (nuc. 8) | | | | 1 | | | | 2 | 1 | | | | | | | 1 | |
| C del. (nuc. 12) | | 2 | | 4 | | | | 2 | | | | | | | | | |
| A del. (nuc. 13) | 1 | | | | | | | | | | | | | | | | |
| C del. (nuc. 14) | | | | | 1 | 1 | 1 | 1 | | | | 1 | | | | | |
| A del. (nuc. 16) | | | | | | | 1 | | | | | | | 1 | | | |
| C del. (nuc. 17) | | | | | | | | | | | | | 3 | | | | |
| T del. (nuc. 18) | | 1 | | 2 | | | | 1 | | | | | | | | | |
| C del. (nuc. 19) | | | | | | | | | | | | 1 | | 1 | | | |
| A del. (nuc. 20) | | | | 7 | 3 | 1 | 1 | 3 | | 2 | 1 | | 2 | | 1 | | |
| C del. (nuc. 21) | | | | | | 1 | | 2 | | | 1 | | 1 | | | | |
| A del. (nuc. 22) | | | | | | | 4 | 2 | 1 | | | | | | | | |
| C del. (nuc. 23) | | | | 2 | | 2 | 1 | | | | | | | | 1 | | |
| > -1 del | 2 | | | 1 | 1 | | | 1 | | 2 | | | | 1 | 0 | 0 | 0 |
| +2 add. | 4 | | | 3 | 3 | | 3 | 3 | 1 | 4 | 3 | 0 | | 0 | 3 | 2 | 2 |
| > +2 add. | 3 | | | 4 | 3 | 1 | 1 | 2 | 1 | 0 | 2 | 3 | | 1 | 0 | 3 | 1 |
| subst. | | | | 5 | | | 3 | 13 | 3 | 3 | 5 | 0 | 3 | 5 | 4 | 6 | 4 |
| no *leuD* mut. | 13 | 3 | 1 | 18 | 14 | 0 | 5 | 12 | 18 | 16 | 11 | 17 | 6 | 10 | 9 | 6 | 11 |
| Total | 41 | 26 | 12 | 82 | 28 | 32 | 42 | 49 | 40 | 34 | 33 | 29 | 35 | 24 | 24 | 24 | 18 |

TableS4: Main TLS polymerases dependent *rpoB* mutations found in Mtb clinical isolates

| *M. smegmatis* | | | | | *E. coli* | Mtb | | WHO report (rif[R] clin. isol.) | | |
|---|---|---|---|---|---|---|---|---|---|---|
| amino acids | mut. (aa) | mut. (codon) | freq. empty | freq. *dinB1*[Mtb+5aa] OE | aa | aa | codon | mut. | freq. | total nbr. |
| His442(H) | His(H)>Arg(R) | CAC>CGC | 1.19 | 33.19 | His526 | His445 | CAC | H445R | 0.8% | 79 |

| *M. smegmatis* | | | | | *E. coli* | Mtb | | WHO report (rif[R] clin. isol.) | | |
|---|---|---|---|---|---|---|---|---|---|---|
| amino acids | mut. (aa) | mut. (codon) | freq. Δ*dnaE2* $H_2O_2$ | freq. WT $H_2O_2$ | aa | aa | codon | mut. | freq. | total nbr. |
| Ser447(S) | Ser(S)>Leu(L) | TCG>TTG | 4.33 | 11.98 | Ser531 | Ser450 | TCG | S450L | 66.2% | 6536 |
| Ser438(S) | Ser(S)>Leu(L) | TCG>TTG | 0.43 | 8.99 | Ser522 | Ser441 | TCG | S441L | 0.3% | 26 |
| his442(H) | His(H)>Tyr(Y) | CAC>TAC | 1.73 | 7.49 | His526 | His445 | CAC | H445Y | 3.5% | 347 |
| his442(H) | His(H)>Asp(D) | CAC>GAC | 1.73 | 5.99 | His526 | His445 | CAC | H445D | 2.9% | 288 |
| Asn435(N) | Asn(N)>Lys(K) | AAC>AAG | 0.00 | 4.49 | Asn519 | Asn438 | AAC | N438K | ND | ND |
| Leu427(L) | Leu(L)>Pro(P) | CTG>CCT | 1.30 | 3.00 | Leu511 | Leu430 | CTG | L430P | 1.1% | 106 |
| Gln429(Q) | Gln(Q)>Leu(L) | CAG>CTG | 0.00 | 3.00 | Gln513 | Gln432 | CAA | Q432L | ND | ND |
| Ser438(S) | Ser(S)>Trp(W) | TCG>TGG | 0.00 | 3.00 | Ser522 | Ser441 | TCG | S441W | ND | ND |
| Arg445(S) | Arg(R)>Cys(C) | CGT>TGT | 0.43 | 3.00 | Arg529 | Arg448 | CGA | R448C | ND | ND |

Freq.: absolute frequency of *rpoB* mutations, expressed in number of mutation per $10^8$ CFU found in indicated strains (also shown in Figures 2D and 3C).
a.a.: *rpoB* amino acids of *M. smegmatis* found in rif[R] colonies and corresponding amino acids in the *E. coli* and Mtb *rpoB* gene.
WHO report: frequencies (freq.) and number (total nbr.) of Mtb rif[R] clinical isolates in which listed *rpoB* mutations have been detected among around 10 000 sequenced Mtb rif[R] clinical isolates. The data are published in the WHO mutations catalogue, 2021.
The mutations listed in this table are the mutations detected at a frequency ≥ $3/10^8$ CFU in *M. smegmatis* the condition in which the TLS polymerase is expressed.
